# *In situ*, fluorescence lifetime-based measurements of cell membrane micromechanics

**DOI:** 10.1101/694620

**Authors:** S Son, HS Muddana, C Huang, S Zhang, PJ Butler

## Abstract

Microscopic *in situ* measurements of the mechanical properties of lipid bilayers were derived from the mean and variance of the fluorescence lifetime distributions of 1’-dioctadecyl-3,3,3’3’-tetramethylindocarbocyanine perchlorate (DiI). In this method, DiI, incorporated into membranes, acts as a membrane-targeted molecular rotor whose fluorescence lifetime is sensitive to local lipid viscosity. A new model was developed in which changes in area per lipid were derived from the first and second moments of a stretched exponential distribution of fluorescence lifetimes of DiI, which were subsequently used to compute mean area per lipid and its variance, quantities directly related to bilayer compressibility and bending moduli. This method enabled molecular scale assays of surface micromechanics of membrane-bound entities, such as nanoliposomes and human red blood cells.

**STATEMENT OF SIGNIFICANCE:** Despite the progress in cell deformability studies, and in understanding mechanical properties of purified lipid bilayers, there has not, to date, been a method to measure the mechanics of the lipid bilayer in cells in situ. The current manuscript describes such a method. Using a fluorescent molecular rotor, DiI, embedded in the membrane, along with time resolved fluorescence, we directly measure area per lipid, and its temporal and spatial variance, properties directly related to bilayer mechanical moduli. Such a method will allow investigators to start exploring the relationship between lipid bilayer mechanics and cellular health and disease.

---

The elastic moduli of membranes are fundamental to the understanding of cellular responses to mechanical stimuli (reviewed in (1)), the sensitivity of ion channels to membrane stretch (reviewed in (2)), responses of cilia to fluid flow (reviewed in (3)) and mechanobiological phenomena responsible for health and disease (reviewed in (4)). Derived from area per lipid (APL), these moduli provide input to models that predict membrane bending fluctuations, by which cells sense their surroundings (5), and hydrophobic matching between lipids and proteins, key constants governing lipid phase separation and protein modulation by lipids (6, 7). Thus, APL and associated moduli are fundamental mechanistic links between lipid membrane composition and membrane-related biological functions of cells.

Current methods to measure membrane APL include nuclear magnetic resonance (NMR) spectroscopy, X-ray diffraction and neutron diffraction (listed in Table S2) that derive APL from measurements of thermally induced displacement of stacked bilayers from equilibrium positions, acyl chain conformations, bilayer thickness, or other membrane structural parameters. Because these techniques require that lipids are isolated, they are not suitable for *in situ* measurements in cells. Alternatively, cellular membrane moduli have been assessed using high-resolution optical registration of thermally induced surface fluctuations (reviewed in (8)), micromechanical perturbation by a pipette (9), or atomic force microscopes (10). These methods determine moduli directly using a relationship between membrane deformation and applied force. However, these methods do not provide APL, do not probe the bilayer exclusively, and often require cells to be isolated and attached to surfaces, which may alter the very mechanical properties being assessed.

Recently, Colom and colleagues developed a fluorescence lifetime-based method in which an exogenous lipophilic molecular tension sensor was introduced to live cells (11). Their planarizable push-pull probe revealed a fluorescence lifetime that decreased linearly with increases in membrane tension. Although not measured, their method could potentially be used to measure bilayer mechanical properties provided that the relationship between tension and areal strain was known. In our studies, we seek to measure mechanical properties directly. In this work we present a new time-resolved fluorescence microscopy-based technique that enables the measurement *in situ* of cell membrane micromechanics in a noninvasive manner. We relate the mean fluorescence lifetime of lipid bilayer-bound 1’-dioctadecyl-3,3,3’3’-tetramethylindocarbocyanine perchlorate (DiI) from a stretched exponential distribution of lifetimes to local molecular free volume and, hence, area per lipid (APL), and the variance of the APL distribution to compressibility and bending moduli of the membrane.

The fluorescence lifetime of DiI, an indocarbocyanine dye that belongs to the cyanine dye family, is strongly influenced by the *trans-cis* photoisomerization from the first excited state, caused by carbon-carbon bond rotation about the central methine bridge (12, 13) (Fig. 1). Because this rotation depends on the local viscosity, the lifetime can be shown to be a function of local molecular area. In our previous molecular simulation of DiI and lipids, we showed that DiI intercalated between lipids just below the lipid headgroup, making fluorescence lifetime dependent on the viscosity at the headgroup-acyl chain interface (14). In Supplementary Note 1, we show that the average fluorescence lifetime of DiI is related to APL according to:

**Figure 1.**
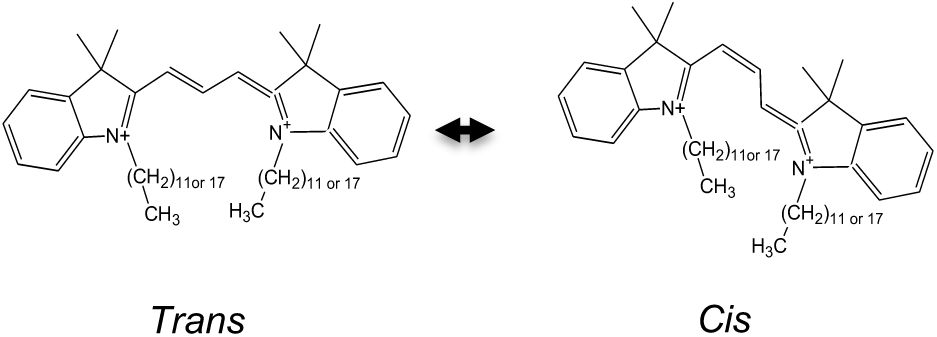
Structure of DiI and trans-cis isomerization of the chromophore.

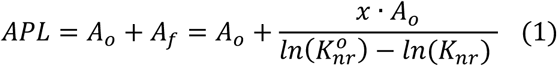

where *A*_*f*_ is the free area, *A*_*o*_ is the hardcore van der Waals area (0.42 nm^2^ for phosphatidylcholine (PC)(15), 0.34 nm^2^ for sphingomyelin (SM)(16), and 0.30 nm^2^ for cholesterol(17), 0.38 nm^2^ for cells (65% of PC, 10% of SM, and 25% of cholesterol (18))), *x* is a constant determined in solution experiments (0.37 Fig. S3A), *ln*(*K*_*nr*_) = − 1.064 * *In*(⟨*τ*⟩) − 1.4403 (Fig. S3C), and 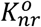 is the non-radiative decay rate in low viscosity.

The lifetime of DiI is generally characterized by a double exponential decay function (19). However, in order to apply this method to cells, the membranes of which are characterized by a heterogenous lipid composition, fluorescence lifetime was captured using a stretched exponential distribution function (20). Curve fitting was done using Fluofit software (PicoQuant Gmbh, Berlin, Germany) by a process of iterative reconvolution given by,

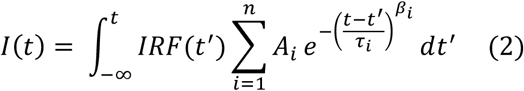

where *I(t)* is the experimental decay function, *IRF(t)* is the instrument response function, *A*_*i*_ is the amplitude of *i*^*th*^ component, *τ*_*i*_ is the lifetime of *i*^*th*^ component, *β*_*i*_ is the distribution parameter of *i*^*th*^ component. Usually 0 < *ß* < 1. *ß* =1 is equivalent to an exponential decay. The mean ⟨*τ*⟩ and variance 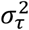 of lifetimes were calculated using the following equations:

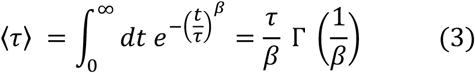

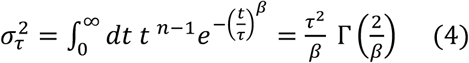

where Γ is the gamma function.

To calculate the variance of APL, we used the delta method after further expansion of Eq. 1 in a Taylor series about the mean of DiI lifetime ⟨*τ*⟩. The series was truncated to the first-order because the function is an approximately linear relationship between observed lifetime range and APL (Fig. S4). The first-order ap-proximation of variance (_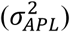_) of APL is:

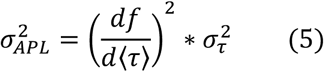

It is known that the thermally driven areal fluctuations of lipids around a mean area can provide the area compressibility modulus, *K*_*A*_, of the lipid bilayer according to(17):

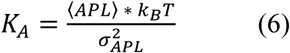

where *k*_*B*_ is Boltzmann constant, *T* is temperature, and 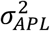 is the variance of the APL distribution. Assuming a polymer brush model for the lipid bilayer (21), the membrane bending modulus *κ*_*c*_can be calculated from *K*_*A*_ by,

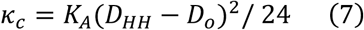

where *D*_*HH*_ is the head-to-head thickness of the bilayer (Table S3) and *D*_*o*_ = 1.0 nm is double the distance between the end of the hydrophobic core and center of the bilayer headgroup (21).

We first measured the fluorescence lifetime of DiI embedded in ∼100 nm nanoliposomes prepared from both saturated and unsaturated lipids at various temperatures ranging from 24 °C to 42 °C (Fig. 2A) using TCSPC (22) and converted lifetimes to APL using Eq. 1. APL values of all lipids increased with increases in temperature. Thermal expansion coefficients α_A_ of different compositions in liquid phase were the slope of this curve at 30°C, and ranged from 0.0071 °C^-1^ to 0.0121 °C^-1^, depended on chain length and the degree of unsaturation (Table S4), and were slightly higher than some literature values (23). Increasing chain length in saturated lipids resulted in a decrease in thermal expansion coefficients. Compressibility modulus, *K*_*A*_, computed for nanoliposomes from various lipids at temperatures ranging from 24 °C to 42 °C indicated dependence on temperature of APL, *K*_*A*_, and *κ*_*c*_ (Figs. 2 A-C), similar to the dependence on temperature of *K*_*A*_ for DOPC (24).

**Figure 2.**
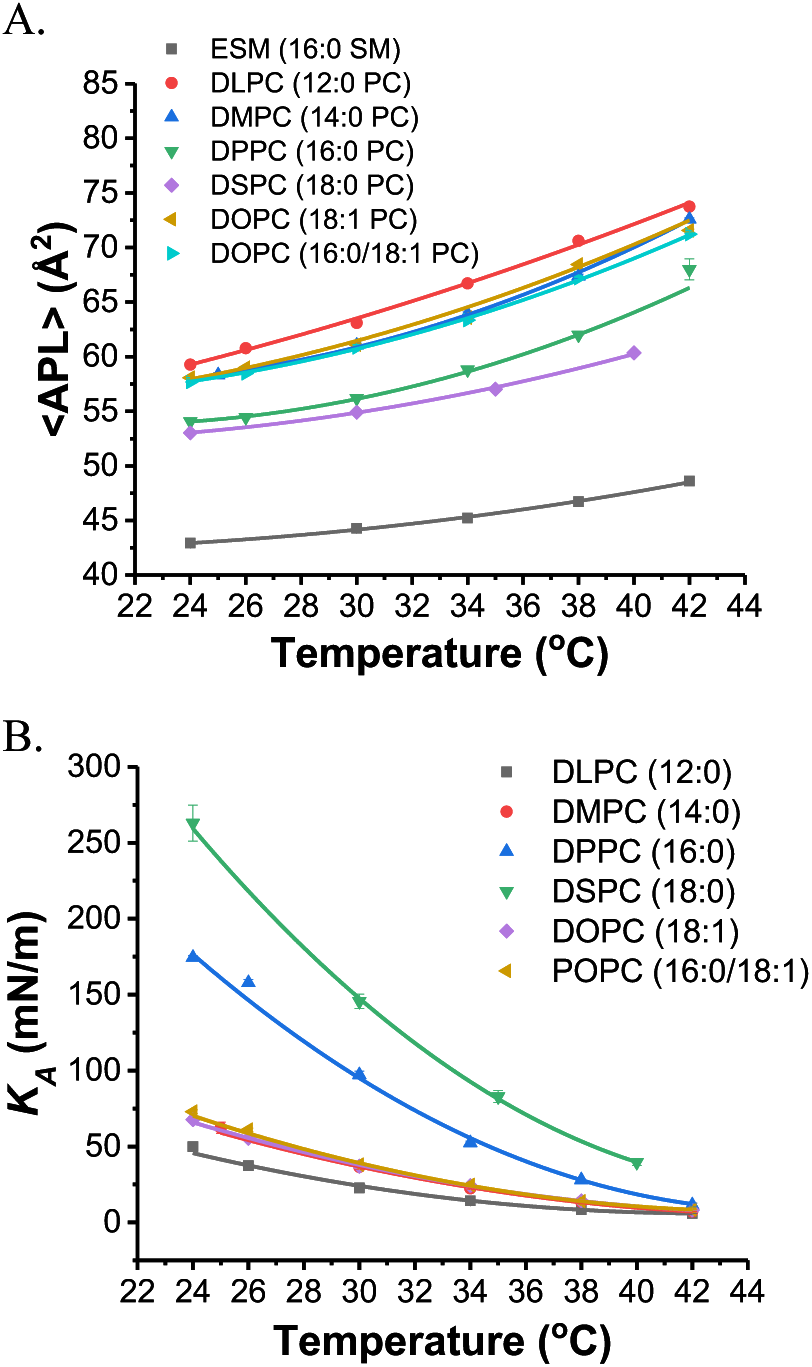
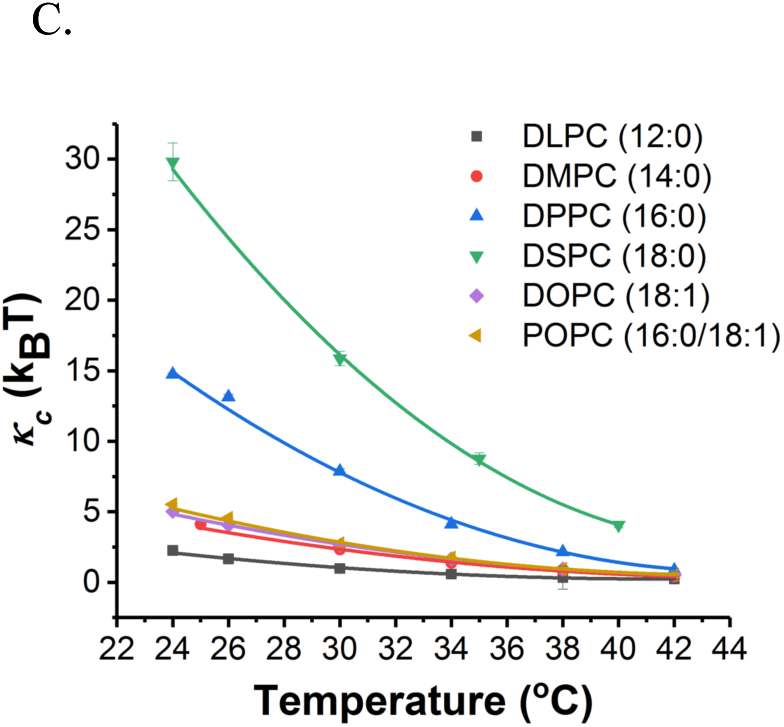
(A) Area per lipid (APL), (B) compressibility moduli (*K*_*A*_), and (C) bending moduli (*κ*_*c*_) for the various lipids of nanoliposomes determined by assessing the fluorescence lifetime distribution through a range of temperatures between 24 °C and 42 °C for different compositions. All error bars are the standard error of the mean (S.E.M) of five measurements.

APL and moduli were also sensitive to the lipid type. The APL values decreased with increasing lipid acyl chain length at constant temperature (Fig. S5). The gel phase lipid, DSPC (18:0), exhibited the highest *K*_*A*_ of 263 mN/m and *κ*_*c*_ of 29.8 k_B_T at 24 °C and a successive decrease in chain length of 2 carbons corresponded to lower compressibility moduli of 174, 62, and 50 mN/m and bending moduli of 14.8, 4.1, and 2.3 k_B_T for DPPC (16:0), DMPC (14:0), and DLPC (12:0), respectively. To apply membrane stress at constant temperature, we increased osmotic pressure of nanoliposomes by adjusting solute concentration gradients across the membrane. This procedure caused increases in APL and the slope of APL versus osmotic pressure for liquid phase lipids was steeper than for gel phase lipids (Fig. S6).

Our goal was to develop a method that could be used, *in situ*, on intact membranes of live cells. Therefore, we examined whether membrane moduli could be measured in human red blood cells. The value of *κ*_*c*_ of human red blood cells (hRBCs) at 24 °C was 2.57 k_B_T (shown in Table 1) and is comparable to available measurements done with diffraction phase microscopy (25) but lower than the value from weak optical tweezers (26). Differences may illustrate effects of cell adhesion to surfaces, limitations of the polymer brush model for membranes containing proteins, or the fact that our probe measures lipid environments directly, irrespective of membrane-spectrin interaction. Increases in temperature to a physiological temperature led to increases in APL and decreases in values of *K*_*A*_ and *κ*_*c*_, consistent with the results from nanoliposomes. Regarding the role of ATP in RBCs, some research demonstrates that the value of *κ*_*c*_ of membranes increases after ATP-depletion because of the enhanced association of the membrane to the spectrin network upon ATP depletion (26). Interestingly, our technique revealed that ATP-depletion caused hRBC shape transitions from discocyte to echinocyte and ovalocyte with accompanying increases in area per lipid and decreases in the values of *K*_*A*_ and *κ*_*c*_, a phenomenon that mirrored results from echinocytes and ovalocytes without drug treatments (Fig. S7 and Table S5). Although requiring further experiments, we hypothesize that the shape of RBCs is likely controlled by the cytoskeleton, and that changes in shape indirectly change the mechanical properties of the lipid bilayer. Further, probing the mechanical properties of the lipid membrane is a convenient way to assess the subtle attachments of the membrane to the cytoskeleton, which are metabolically controlled. For example, enhanced spicules of RBC membrane upon the loss of ATP increased the free area of the membrane and eventually decreased the energy requirement to bend the membrane, as evidenced by the lower bending modulus measured by our technique.

**Table 1.**
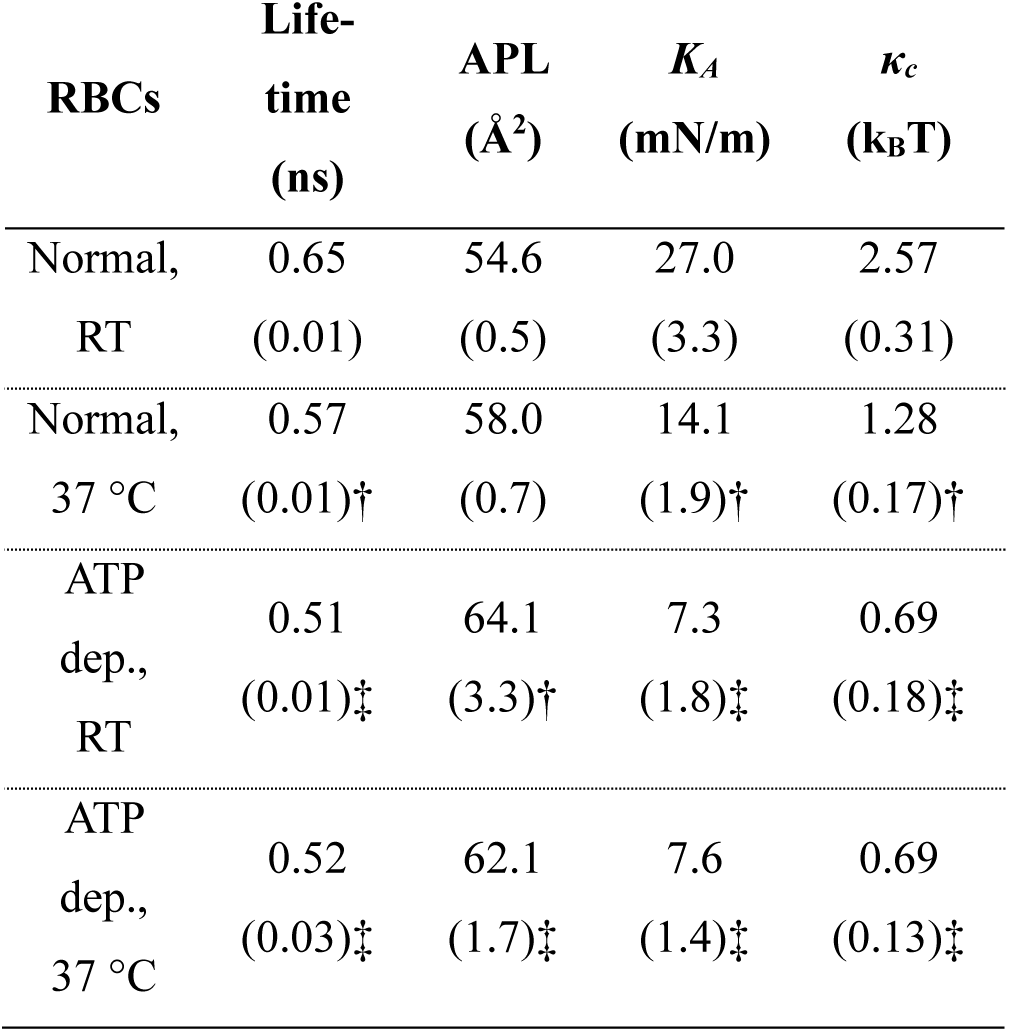
The values of APL, *K*_*A*_ and *κ*_*c*_ in human red blood cells (hRBCs). ATP was depleted with Inosine and iodoacetamide. The experiments conducted at were room temperature (RT = 24 °C) and physiological temperature (37 °C) as indicated. Data represent the mean (S.E.M) of 9-10 cells. Statistical analyses for APL were done by Kruskal-Wallis test followed by Dunne’s multiple comparison test one-way and others were completed by ANOVA followed by Holm-Sidak’s multiple comparisons test; *:p<0.05, †:p<0.01, and ‡:p<0.0001. All significance values are relative to normal control at RT.

We predict that this technique will accelerate our fundamental understanding of the role of membrane mechanics in a wide range of physiological cellular processes. Since the cellular membrane has high compositional and structural complexity, accurate measurements of areal changes in cell membranes would open up several new avenues for understanding the role of hydrophobic mismatch, fluidity, stretch, and curvature in regulating membrane protein activity in live cells (27). Since changes in lipid packing under the influence of an externally applied stimulus are rapid and transient, such a method may have the necessary spatial and temporal precision at the sub-micron and sub-second levels(28) to assess these mechanobiological aspects of the cell membrane. Such rapid and automated measurements of membrane mechanical properties might aid in the optimization of liposome formulations for drug delivery, help in the understanding of mechanical barriers to nanoparticle uptake by cells (29) or become an assay for diseases characterized by changes in membrane biophysics such as modification of the RBC membrane by malarial parasites (30).

## SUPPORTING MATERIAL

Material and methods and Supplementary note 1 (PDF)

## AUTHOR CONTRIBUTIONS

SS and HSM conducted experiments and contributed to writing the manuscript, CH and SZ assisted in interpreting bending moduli from fluorescence lifetime distributions, PJB conceived the project, assisted in experiments and interpretation of data, and helped write and edit the manuscript.

## ACKNOWLEDGMENT

PJB Acknowledges funding from the National Science Foundation (NSF CMMI 1334847).

## References

1. Bao, G., and S. Suresh. 2003. Cell and molecular mechanics of biological materials. Nat. Mater. 2: 715–725.

2. Kung, C. 2005. A possible unifying principle for mechanosensation. Nature. 436: 647–54.

3. Singla, V., and J. Reiter. 2006. The Primary Cilium as the Cell’s Antenna: Signaling at a Sensory Organelle. Science (80-.). 313: 629–633.

4. Jaalouk, D.E., and J. Lammerding. 2009. Mechanotransduction gone awry. Nat. Rev. Mol. Cell Biol. 10: 63–73.

5. Waheed, Q., and O. Edholm. 2009. Undulation Contributions to the Area Compressibility in Lipid Bilayer Simulations. Biophys. J. 97: 2754–2760.

6. Mouritsen, O.G., and M. Bloom. 1993. Models of lipid-protein interactions in membranes. Annu. Rev. Biophys. Biomol. Struct. 22: 145–71.

7. Son, S., G.J. Moroney, and P.J. Butler. 2017. beta1-Integrin-Mediated Adhesion Is Lipid-Bilayer Dependent. Biophys. J. 113: 1080–1092.

8. Monzel, C., and K. Sengupta. 2016. Measuring shape fluctuations in biological membranes. J. Phys. D. Appl. Phys. 49: 243002.

9. Hochmuth, R.M. 2000. Micropipette aspiration of living cells. J. Biomech. 33: 15–22.

10. Dulinska, I., M. Targosz, W. Strojny, M. Lekka, P. Czuba, W. Balwierz, and M. Szymonski. 2006. Stiffness of normal and pathological erythrocytes studied by means of atomic force microscopy. J. Biochem. Biophys. Methods. 66: 1–11.

11. Colom, A., E. Derivery, S. Soleimanpour, C. Tomba, M.D. Molin, N. Sakai, M. González-Gaitán, S. Matile, and A. Roux. 2018. A fluorescent membrane tension probe. Nat. Chem..

12. Widengren, J., and P. Schwille. 2000. Characterization of Photoinduced Isomerization and Back-Isomerization of the Cyanine Dye Cy5 by Fluorescence Correlation Spectroscopy.: 6416–6428.

13. Sanborn, M.E., B.K. Connolly, K. Gurunathan, and M. Levitus. 2007. Fluorescence properties and photophysics of the sulfoindocyanine Cy3 linked covalently to DNA. J. Phys. Chem. B. 111: 11064–11074.

14. Muddana, H.S., R.R. Gullapalli, E. Manias, P.J. Butler, and M. E. 2011. Atomistic simulation of lipid and DiI dynamics in membrane bilayers under tension. Phys. Chem. Chem. Phys. 13: 1368–78.

15. Gudmand, M., M. Fidorra, T. Bjørnholm, and T. Heimburg. 2009. Diffusion and partitioning of fluorescent lipid probes in phospholipid monolayers. Biophys. J. 96: 4598–4609.

16. Kupiainen, M., E. Falck, S. Ollila, P. Niemelä, A.A. Gurtovenko, M.T. Hyvönen, M. Patra, M. Karttunen, and I. Vattulainen. 2005. Free volume properties of sphingomyelin, DMPC, DPPC, and PLPC bilayers. J. Comput. Theor. Nanosci. 2: 401–413.

17. Falck, E., M. Patra, M. Karttunen, M.T. Hyvönen, and I. Vattulainen. 2004. Lessons of Slicing Membranes: Interplay of Packing, Free Area, and Lateral Diffusion in Phospholipid/Cholesterol Bilayers. Biophys. J. 87: 1076–1091.

18. van Meer, G., and A.I.P.M. de Kroon. 2011. Lipid map of the mammalian cell. J. Cell Sci. 124: 5–8.

19. Packard, B.S., and D.E. Wolf. 1985. Fluorescence lifetimes of carbocyanine lipid analogs in phospholipid bilayers. Biochemistry. 24: 5176–5181.

20. Lee, K.C.B., J. Siegel, S.E.D. Webb, S. Lévêque-Fort, M.J. Cole, R. Jones, K. Dowling, M.J. Lever, and P.M.W. French. 2001. Application of the Stretched Exponential Function to Fluorescence Lifetime Imaging. Biophys. J. 81: 1265–1274.

21. Rawicz, W., K.C. Olbrich, T. McIntosh, D. Needham, and E. Evans. 2000. Effect of chain length and unsaturation on elasticity of lipid bilayers. Biophys. J. 79: 328–339.

22. Gullapalli, R.R., T. Tabouillot, R. Mathura, J.H. Dangaria, and P.J. Butler. 2015. Integrated multimodal microscopy, time-resolved fluorescence, and optical-trap rheometry: toward single molecule mechanobiology. J. Biomed. Opt. 12: 014012.

23. Kucerka, N., Y. Liu, N. Chu, H.I. Petrache, S. Tristram-Nagle, and J.F. Nagle. 2005. Structure of Fully Hydrated Fluid Phase DMPC and DLPC Lipid Bilayers Using X-Ray Scattering from Oriented Multilamellar Arrays and from Unilamellar Vesicles. Biophys. J. 88: 2626–37.

24. Pan, J., S. Tristram-Nagle, N. Kucerka, and J.F. Nagle. 2008. Temperature Dependence of Structure, Bending Rigidity, and Bilayer Interactions of Dioleoylphosphatidylcholine Bilayers. Biophys. J. 94: 117–124.

25. Park, Y., C.A. Best, K. Badizadegan, R.R. Dasari, M.S. Feld, T. Kuriabova, M.L. Henle, A.J. Levine, and G. Popescu. 2010. Measurement of red blood cell mechanics during morphological changes. Proc. Natl. Acad. Sci. U. S. A. 107: 6731–6.

26. Betz, T., M. Lenz, J.-F. Joanny, and C. Sykes. 2009. ATP-dependent mechanics of red blood cells. Proc. Natl. Acad. Sci. U. S. A. 106: 15320–15325.

27. Lee, A.G. 2003. Lipid-protein interactions in biological membranes: A structural perspective..

28. Tabouillot, T., H.S. Muddana, and P.J. Butler. 2011. Endothelial Cell Membrane Sensitivity to Shear Stress is Lipid Domain Dependent. Cell. Mol. Bioeng. 4: 169–181.

29. Huang, C., P.J. Butler, S. Tong, H.S. Muddana, G. Bao, and S. Zhang. 2013. Substrate stiffness regulates cellular uptake of nanoparticles. Nano Lett. 13: 1611–1615.

30. Callan-Jones, A., O.E. Albarran Arriagada, G. Massiera, V. Lorman, and M. Abkarian. 2012. Red blood cell membrane dynamics during malaria parasite egress. Biophys. J. 103: 2475–2483.

