## Supplementary material for "*In situ*, fluorescence lifetime-based measurements of cell membrane micromechanics": Suplemental Information (from another version of this preprint: version 1, 2019-07-12)

### Experimental Section

#### *DiI and lipids*

1,2-dilauroyl-sn-glycero-3-phosphocholine (DLPC), 1,2-dimyristoyl-sn-glycero-3-phosphocholine (DMPC), 1,2-dipalmitoyl-sn-glycero-3-phosphocholine (DPPC), 1,2-distearoyl-sn-glycerol-3-phosphocholine (DSPC), 1,2-dioleoyl-sn-glycero-3-phosphocholine (DOPC), 1-palmitoyl-2-oleoyl-sn-glycero-3-phosphocholine (POPC) were purchased from Avanti polar lipids, Inc. Fluorescent probes 1,1'-didodecyl-3,3,3',3'-tretramethylindocarbocyanine perchlorate (DiI-C12; $\lambda ex = 549$ nm; $\lambda em = 565$ nm) was purchased from Invitrogen, Inc. All the materials were obtained at their highest purity available and used without further purification.

#### *Absorption, emission spectra, and quantum yield measurements*

Absorption and emission spectra of DiI were measured in ethanol/glycerol mixture. Absorption spectra were obtained using the Agilent 8453 UV-Visible spectrophotometer (Agilent Technologies, Inc.) with spectral bandwidth of 1 nm. Emission spectra were obtained using RF-5301PC spectrofluorometer (Shimadzu, Inc.), with the spectral bandwidth of 1.5 nm. The emission spectra were collected at an excitation wavelength of 532 nm. Viscosities of the different solvents were measured using a cone-plate viscometer (RotoVisco 1, Rheology Solutions, Inc.). Refractive indices of binary mixtures of ethanol and glycerol were determined from the measured viscosity using the empirical relationship reported by Alkindi et al. (1). Quantum yield ($\Phi_S$) was calculated with reference to Rhodamine 6G ($\Phi_R = 0.95$ in ethanol), using the following equation (2):

$$\Phi_S = \Phi_R \frac{IE_S}{IE_R} \frac{A_R}{A_S} \frac{(n_S)^2}{(n_R)^2} \quad (6)$$

where the subscripts *R* and *S* refer to rhodamine and the unknown sample, respectively, *IE* is the integrated emission intensity, *A* is absorbance at 532 nm, and *n* is the refractive index.

*Fluorescence lifetime and fluorescence correlation spectroscopy (FCS)*

Fluorescence lifetime measurements were conducted using a time-correlated single photon counting (TCSPC) instrument developed in our lab (3). Briefly, the instrument was based on an Olympus IX71 inverted microscope (Tokyo, Japan) and a water-cooled 80 MHz, 5.4 ps pulse width, 75 mW power, pulsed solid state laser (λ = 532 nm, High-Q Laser, Hohenems, Austria) was used as an excitation source. The laser output was coupled to an optical fiber and was cleaned by an excitation filter and expanded to slightly underfill the back aperture of the objective. The laser power as measured at the back aperture of the objective was adjusted to a value between 50 and 100 µW for nanoliposome or 30 µW for cells to avoid photobleaching and triplet state formation. Laser light was focused in the sample using a 60X water immersion objective (UPLAPO60XW, Olympus). Fluorescence emission from the sample was passed through a set of high-quality emission filters, and a polarizer fixed at the magic angle of 54.7$^{o}$, before focusing onto a 50 µm/0.22NA optical fiber that served as the confocal pinhole. The optical fiber was coupled to a GaAsP photomultiplier tube (H7422-40P, Hamamatsu, Tokyo, Japan) with a peak quantum efficiency of 40% at 580 nm wavelength. Photon arrival times were recorded using the SPC-630 module from Becker and Hickl (Berlin, Germany). The instrument response function (IRF) was collected using a solution of 200 mM rhodamine 6G or rhodamine B in methanol (4) prior to the experiment. Histograms of microtimes from TCSPC were fit using iterative reconvolution of an IRF with a stretched exponential distribution function using Fluofit software (PicoQuant Gmbh, Berlin, Germany). The quality of the fitted curves was evaluated based on $\chi^2$ values.

Diffusion coefficients of liposome were determined by fitting the autocorrelation curves to single-component three-dimensional diffusion model. The quality of the fitted curves was evaluated based on chisquare ($\chi^2$) values. Radius of the observation volume determined using the known diffusion coefficient of Rhodamine 6G (D = 2.8 x $10^{-10}$ $m^2/s$ in water) or 50 nm fluorescence bead was ~400 nm. Radii of particles were determined from the diffusion coefficient using the Stokes-Einstein relationship. Positioning of the membrane (model and cell membranes) along the optical axis was achieved by maximizing the fluorescence intensity while performing an intensity scan.

*Preparation of nanoliposomes with DiI*

Lipid/dye mixture solutions were prepared to a concentration of 0.25 mg/ml with the dye concentration of ~10 nM for FCS. Nanoliposomes were prepared using the extrusion method (5) as follows. 1 mg of lipid in chloroform was deposited in a round bottom flask and dried under argon for 5 minutes and vacuum dried for an hour. 1 ml of water was then introduced into the flask and maintained at a temperature of 70 °C. The flask was frequently agitated for up to an hour. This liposome suspension was then extruded several times ( > 15) through a polycarbonate membrane filter of 100 nm pore size using the mini-extruder from Avanti lipids, Inc. This typically resulted in a homogeneous liposome sample with sizes on the order of 100 to 150 nm diameter, as verified by dynamic light scattering and fluorescence correlation spectroscopy.

To induce osmotic pressure across the lipid bilayer, nanoliposomes prepared in 500 mM NaCl solution were dispersed in diluted concentrations of solute to induce solute concentration gradients across the membrane. The osmolality was measured by a vapor pressure osmometer (5500, Wescor).

*Preparation of hRBCs and DiI staining*

Human RBCs were freshly prepared before each experiment by finger pricking (IRB #: STUDY00002505). Fifteen microliters of blood were diluted in 1000 µL of PBS containing 0.1 % bovine serum albumin (BSA). RBCs washed three times by centrifugation (2 min, 6000 rpm) and cells were diluted 1:100 for DiI staining. Cells were stained with 100 nM DiI for 3 min and washed four times by centrifugation. To maintain the medium temperature at 37°C during experiments, cells were plated into a temperature chamber (Delta T4 Culture Dish, Bioptechs, PA) and the temperature was controlled with a temperature controller (Delta T Culture Dish Controller, Bioptechs, PA). Before seeding cells, all surfaces of chamber were coated with 5 % BSA for 30 min.

For ATP depletion, hRBCs were incubated in a PBS solution without glucose, in the presence of 10 mM inosine (Sigma-Aldrich) and 6 mM iodoaceamide (Sigma-Aldrich) for 3h at 37˚C. After ATP depletion, cells were returned to PBS containing 0.1 % BSA.

**Note 1. Fluorescence lifetime of DiI can be converted to free-volume in solution**

The fluorescence lifetime of DiI, an indocarbocyanine dye that belongs to the cyanine dye family, is determined by its radiative and non-radiative decay rates. In general, the radiative decay rate is affected by the dielectric constant (or refractive index) of the medium (6). Moreover, it is often independent of the excitation wavelength and the emission spectrum, since the vibrational relaxation occurs very fast (~$10^{-12}$ s) in solution and the emission arises from the lowest vibrational level of the excited state. Therefore, the radiative rate is often considered a constant. On the other hand, non-radiative decay is associated with energy dissipation arising from dynamic structural alterations in the molecule. The non-radiative decay rate of cyanines is strongly influenced by the *trans-cis* photoisomerization and this process is schematically represented in Fig. S1, adopted from potential energy diagram for cyanine photoisomerization proposed by Levitus.(7).

The *trans* conformation is the most preferred conformation in the ground state for cyanine dyes. The molecule in the *trans* excited state can deactivate to the *trans* ground state through emission of a photon (i.e. fluorescence). When the motion of dye is restricted, the brightness of dye increases (8). Alternatively, it can undergo isomerization to the *cis* conformation. While the exact nature of the intermediate conformation is not well understood, it is thought that the molecule goes through a partially twisted intermediate state that is very unstable (9). In the partially twisted state, the molecule can deactivate quickly in a non-radiative fashion to the *trans* or *cis* ground states. Then, the photoisomer goes thorough the thermal back isomerization reaction to return the stable all-trans form.

The effective rate of this deactivation depends on the activation barrier associated with partial twisting of the central methine bridge. The activation barrier for torsional rotation of the molecule is thus determined by the steric constraints, in other words microviscosity. Increasing the viscosity of the solvent increases the activation barrier, thus reducing the rate of *trans-cis* isomerization. Because photoisomerization competes with fluorescence, most cyanine dyes exhibit very low quantum yields when in environments allowing substantial intramolecular rotation(10). Restriction of intramolecular rotation results in increased quantum efficiency and fluorescence lifetime, a phenomenon used to detect when cyanine dyes bind to DNA or RNA.[3] Deactivation of the excited state can also occur through intersystem crossing or internal conversion, however, the decay rates associated with these pathways have been shown to be negligible.

Previously, Loutfy and Arnold (1982)(11) have determined the effect of viscosity and temperature on the torsional relaxation of several molecular rotors. According to their theory, the torsional relaxation rate is related to the free-volume of the solvent by the following equation:

$$K_{nr} = K_{nr}^{0} \exp\left(-\frac{x \cdot V_0}{V_f}\right) \qquad \text{(S1)}$$

where $K_{nr}$ is the non-radiative decay rate, $K_{nr}^{0}$ is free-rotor reorientation rate, which is obtained in a low viscosity solution, $x$ is a molecule-specific constant, and $V_o$ and $V_f$ refer to the hard-core volume and the free volume, respectively. Combining Eq. S1 with the Doolittle equation (12) for solvent viscosity as a function of free-volume, i.e. $\eta = Aexp(V_0/V_f)$, where $A$ is a constant that depends on the nature of the liquid, gives the relationship between torsional relaxation rate and solvent viscosity, as follows:

$$ln(K_{nr}) = C - x \cdot ln(\eta) \qquad \text{(S2)}$$

where $C$ is an empirical constant that combines all other constants in the equation. To determine whether DiI behaves as a molecular rotor described by Eq. S2, we measured the absorption and emission spectra of

DiI in binary mixtures of ethanol and glycerol at different solvent viscosities, from which we calculated the fluorescence quantum yield (Φ, QE). Emission intensity, QE, and lifetime increased by several fold with an increase in viscosity, within the full range of viscosities studied (1.005 cP to 1413.8 cP) (Table S1). As discussed earlier, increases in viscosity increase the activation barrier for *trans-cis* isomerization, resulting in a decrease in non-radiative decay rate. In order to determine the radiative and non-radiative decay rates, we measured the fluorescence lifetime in the above solutions. Radiative ($K_r$) and non-radiative ($K_{nr}$) decay rates are related to the QE and fluorescence lifetime τ, according to the following equations:

$$K_r = \frac{\Phi}{\tau};\ K_{nr} = \frac{1-\Phi}{\tau} \tag{S3}$$

We also measured the absorption and emission spectra and the fluorescence lifetime of DiI in binary mixtures of ethanol and glycerol (Fig. S2) and all values are listed in Table S1. Fig. S3A is a plot between the measured viscosity and the non-radiative decay rate and the linear relation is consistent with Loutfy's theory. Moreover, fitting the data to Eq. S.2 yields the DiI-specific empirical constant *x* to be 0.37. $ln(\tau)$ of Dil has a linear relationship with $ln(\eta)$ in both binary mixtures (Fig. S3B). Using the definition of fluorescence lifetime, we rewrite Eq. S1 as follows:

$$V_f = \frac{x \cdot V_0}{ln(K_{nr}^0) - ln(K_{nr})} \tag{S4}$$

This equation directly relates free-volume with experimentally measured fluorescence lifetime: $ln(K_{nr}) = -1.064 * ln(\tau) - 1.4403$ (Fig. S3C). Since radiative decay rate was independent of the solvent viscosity and the change in the refractive index due to the change in solvent viscosity is slight, it is reasonable to assume that the radiative rate is constant. Therefore, lifetime mainly depends on the non-radiative rate (Fig. S3C). Because $V_0 = L \cdot A_0$ , $V_f = L \cdot A_f$ where $A_0$ and $A_f$ are the lipid hard core area, and lipid free area, respectively, we can convert volumes to areas and obtain the area-per-lipid $A_L = A_0 + A_f$:

$$A_L = A_o + A_f = A_o + \frac{x \cdot A_o}{ln(K_{nr}^0) - ln\ (K_{nr})} \tag{S5}$$

which is Eq. 1 in the main document.

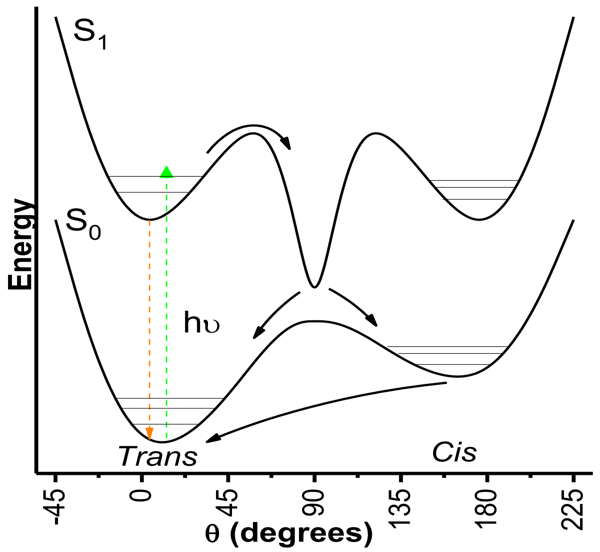


**Figure S1**. Schematic diagram of the energy surface of the ground ($S_0$) and excited ($S_1$) states of DiI. After light absorption, the excited state molecule in S1 completely deactivates via fluorescence emission, internal conversion, and rotation. The partially twisted intermediate molecule deactivates to the all-*trans* ground state or photoisomer (all-*cis* conformation). The photoisomer then goes to the thermodynamically stable all-*trans* isomer without releasing photons. This process involved in *trans-cis* isomerization depends on the viscosity of the medium(7).

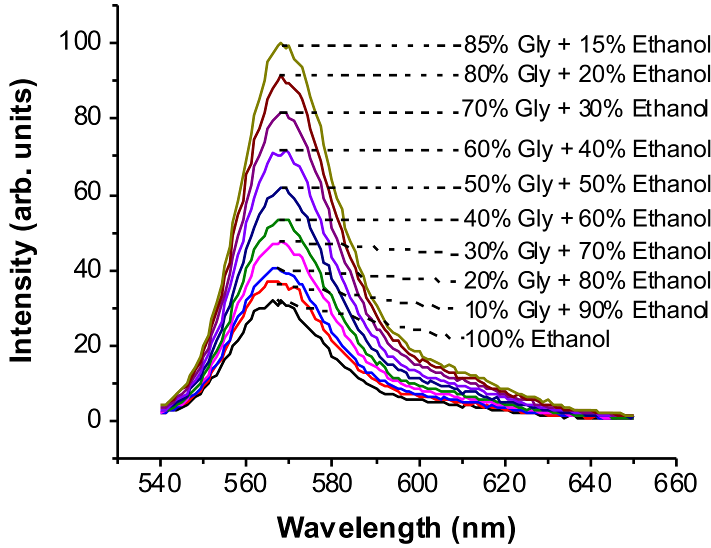

**Figure S2**. Emission spectra of DiI in solutions of different viscosities prepared from binary mixtures of ethanol and glycerol. The corresponding solution viscosities are listed in the Table S1.

A.

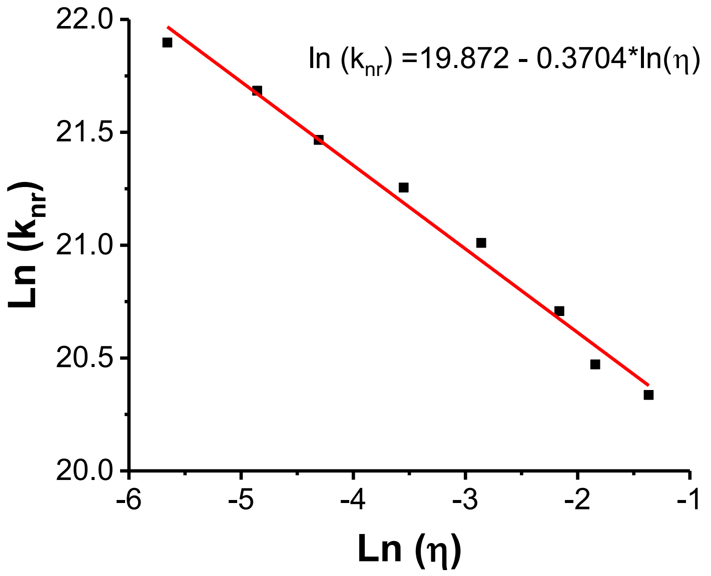


B.

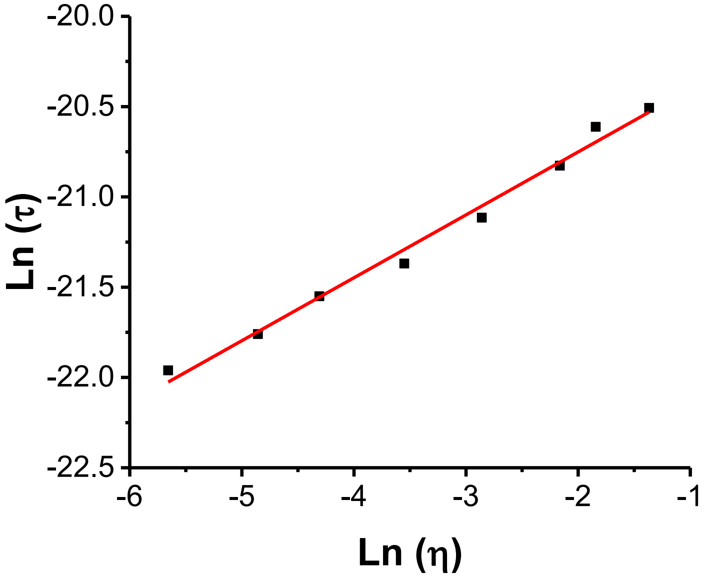

C.

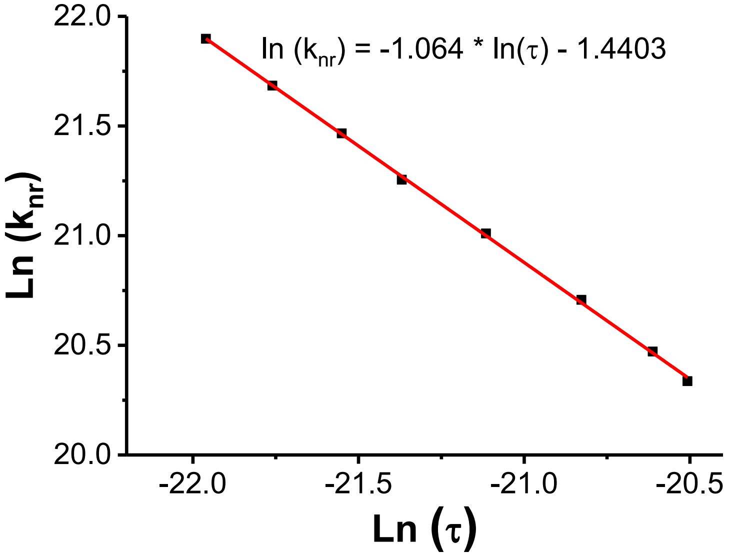


**Figure S3**. Experimental validation of the relationship of (A) non-radiative decay rate (i.e. *trans-cis* photoisomerization rate) and (B) fluorescence lifetime as a function of viscosity. (C) The relationship between non-radiative decay rate (i.e. *trans-cis* photoisomerization rate) and fluorescence lifetime.

**Table S1** Quantum yield (QE), fluorescence lifetime ($\tau$), radiative decay rate ($K_r$), and non-radiative decay rate ($K_{nr}$) of DiI in the various viscosity solutions.

| **S.No.** | **Viscosity (cP)** | **Refractive index** | **Abs (532nm)** | **Integrated Emission** | **QE** | **$\tau$ (ns)** | **$K_r$ ($ns^{-1}$)** | **$K_{nr}$ ($ns^{-1}$)** |
|---|---|---|---|---|---|---|---|---|
| Rhodamine 6G | 1.154 | 1.362 | 0.134 | 31000.115 | 0.950 | 3.667 | 0.259 | 0.014 |
| 20% Gly+80% ethanol | 3.493 | 1.385 | 0.292 | 4164.223 | 0.061 | 0.290 | 0.209 | 3.238 |
| 30% Gly+70% ethanol | 7.781 | 1.400 | 0.287 | 4852.114 | 0.073 | 0.355 | 0.207 | 2.613 |
| 40% Gly+60% ethanol | 13.450 | 1.410 | 0.296 | 5459.882 | 0.081 | 0.437 | 0.186 | 2.102 |
| 50% Gly+50% ethanol | 28.709 | 1.423 | 0.260 | 6282.069 | 0.108 | 0.524 | 0.207 | 1.701 |
| 60% Gly+40% ethanol | 57.294 | 1.434 | 0.331 | 7238.959 | 0.100 | 0.676 | 0.148 | 1.332 |
| 70% Gly+30% ethanol | 115.000 | 1.445 | 0.339 | 8291.309 | 0.113 | 0.901 | 0.126 | 0.984 |
| 80% Gly+20% ethanol | 158.280 | 1.449 | 0.326 | 9209.448 | 0.131 | 1.118 | 0.118 | 0.777 |
| 85% Gly+15% ethanol | 255.040 | 1.455 | 0.303 | 10103.075 | 0.157 | 1.241 | 0.126 | 0.679 |

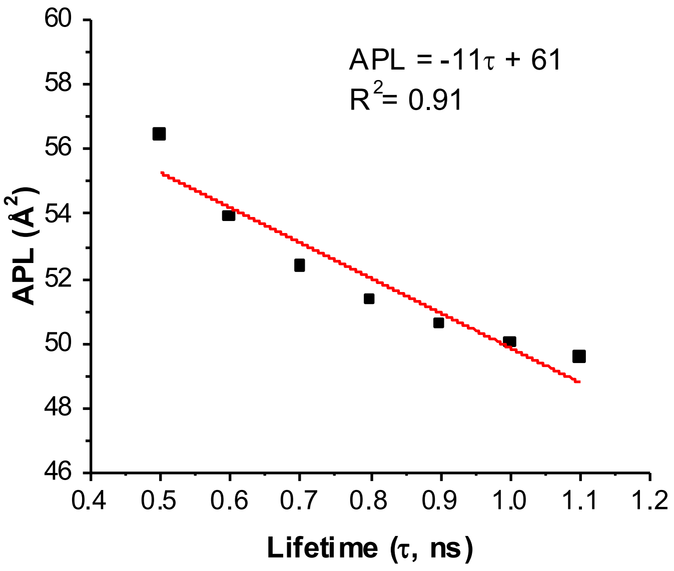


**Figure S4**. The relationship between lifetime, $\tau$, and APL according to Equation S5.

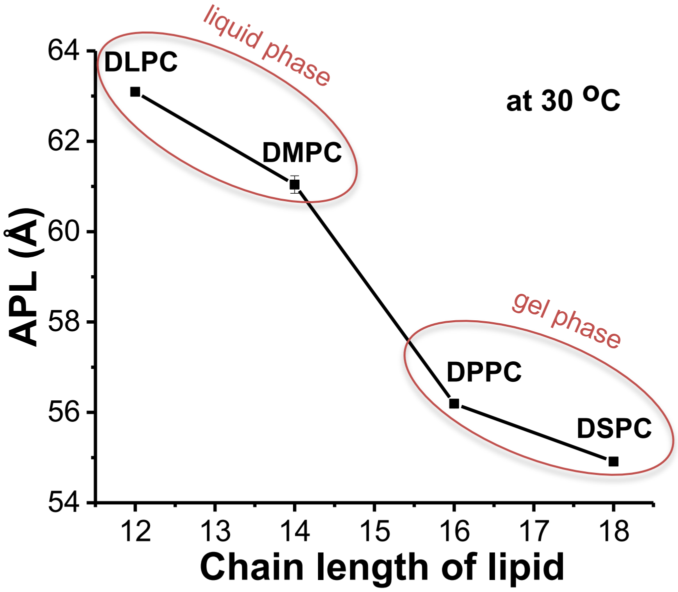


**Figure S5.** The value of APL as a function of chain length of PC at 30 °C in the liquid phase and the gel phase.

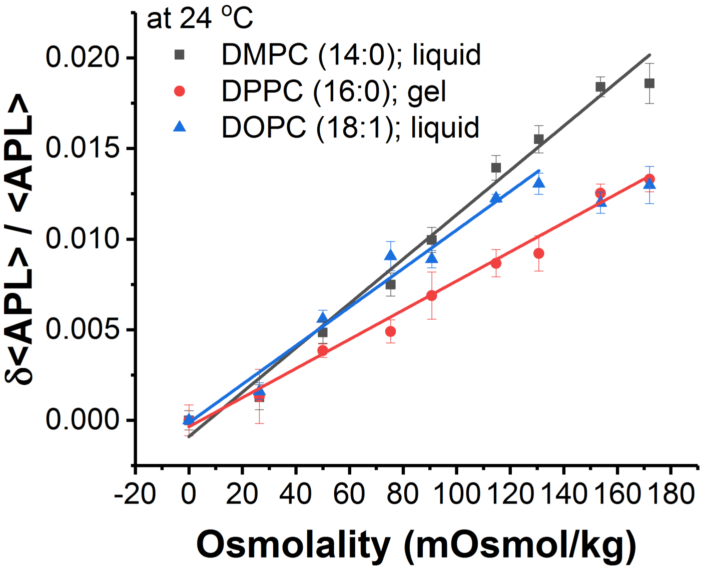


**Figure S6.** Normalized differences in APL of DMPC, DPPC, and DOPC in 500 mM NaCl as a function of changes in osmotic pressure. Osmotic pressure was increased by decreasing external NaCl concentration.

| Lipid | Chain Conf. | Exp APL, Å2 (°C) | Lit APL, Å2 (°C) | Technique |
|---|---|---|---|---|
| ESM | 16:0 (86%) SM | 42.9±0.03 (24) | 47.4 (24) | Langmuir Balances(13) |
| | | 44.3±0.06 (30) | 52.5 (30) | Langmuir Balances(13) |
| DLPC | 12:0 | | 56.7 (10) | NMR(14) |
| | | 59.3±0.16 (30) | 62.6 (30) | NMR(14) |
| | | | 63.2 (30) | X-ray(15) |
| DMPC | 14:0 | 58.3±0.49 (25) | 58.9(24) | Neutron(16) |
| | | | 60.6 (30) | X-ray(15) |
| | | 61.0±0.41 (30) | 59.6 (30) | NMR(14) |
| | | | 59.6 (30) | EDP (17) |
| DPPC | 16:0 | 54.1±0.03 (24) | 47.9 (24) | X-ray (18) |
| | | | 47.3 (25) | X-ray (19) |
| | | 56.2±0.15 (30) | - | |
| DSPC | 18:0 | 53.0±0.20 (24) | 47.3 (25) | X-ray(19) |
| | | 54.9±0.20 (30) | - | |
| DOPC | 18:1 | 58.1±0.23 (24) | 59.4 (23) | X-ray / Neutron(20) |
| | | 61.1±0.25 (30) | 72.5 (30) | EDP (17) |
| | | | 67.4 (30) | X-ray / Neutron(21) |
| POPC | 16:0/18:1 | 57.7±0.14 (24) | 63.0 (24) | Langmuir Balances(22) |
| | | 60.8±0.07 (30) | - | |

**Table S2**. Areas per lipid (Exp APL) calculated from Eq. S5 compared to values obtained by established techniques (Lit APL). NMR is nuclear magnetic resonance spectroscopy and EDP is electron density profile.

| | Temperature (°C) | |
|---|---|---|
| | **30** | **37** |
| DLPC | 3.080 | |
| DMPC | 3.530 | |
| DOPC | 3.670 | |
| POPC | 3.700 | |
| DPPC | 3.853 | |
| DSPC | 4.310 | |
| Cell membrane | | 4.058 |

**Table S3**. Head-to-head thickness ($D_{HH}$ in equation 7) of lipids. The thickness of PC lipids was from Nagle et al. (23) and thickness of cell membranes was obtained from Himhert et al.(24). The temperature dependence of DOPC was used to adjust the thickness of lipid bilayers (25).

| **Lipid** | **Chain Conf.** | **Transition temp (°C)** | **$\alpha_A$=(1/A)(∂A/∂T) at 30 °C** | |
|---|---|---|---|---|
| | | | **Gel phase** | **Liquid phase** |
| DLPC | 12:00 | -2 | | 0.0121 |
| DMPC | 14:00 | 24 | | 0.0103 |
| DPPC | 16:00 | 41 | 0.0094 | |
| DSPC | 18:00 | 55 | 0.0071 | |
| DOPC | 18:01 | -17 | | 0.0098 |
| POPC | 16:0/18:1 | -2 | | 0.0109 |

**Table S4.** Lipid area thermal expansion coefficient, $\alpha_A$

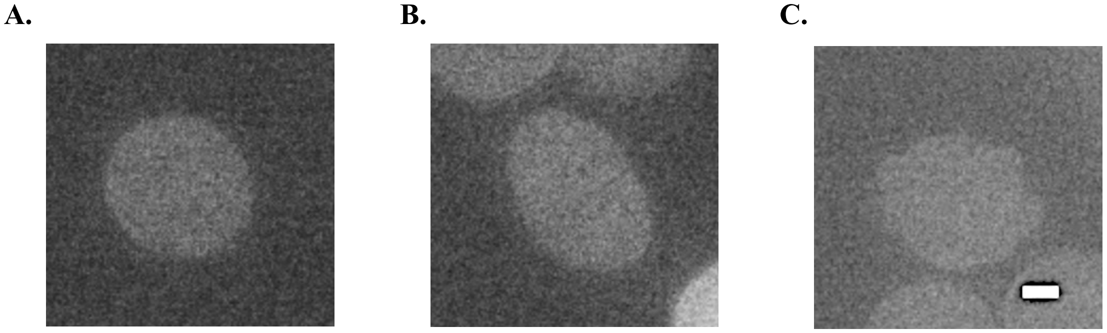


**Figure S7.** Different morphology of hRBC; (A) Normal cell, (B) Ovalocyte, and (C) Echinocyte. Cells were stained with DiI-$C_{12}$ and a fluorescence microscope was used to capture images. Error bar represents 2 μm.

| hRBCs | Lifetime (ns) | APL (Å$^2$) | $K_A$ (mN/m) | $\kappa_c$ ($k_B$T) |
|---|---|---|---|---|
| Normal cells | 0.55 (0.01) | 58.8 (0.5) | 11.4 (1.4) | 1.04 (0.13) |
| Ovalocyte | 0.46 (0.01)† | 67.9 (2.2) | 3.4 (0.7)† | 0.31 (0.06)† |
| Echinocyte | 0.46 (0.02)‡ | 70.0 (3.3)* | 3.9 (0.8)‡ | 0.36 (0.07)‡ |

**Table S5.** The values of lifetime, APL, $K_A$ and $\kappa_c$ in different morphology of hRBCs. The experiments conducted at the 37 °C. Data represent the mean (S.E.M) of 9 -10 cells. Statistical analyses for lifetimes were done by ANOVA followed by Holm-Sidak's multiple comparisons test and others were completed by Kruskal-Wallis test followed by Dunne's multiple comparison test one-way; *:$p<0.05$, †:$p<0.01$, and ‡:$p<0.0001$. All significance values are relative to controls (i.e. normal cells).

**References**

1. Alkindi, A.S., Y.M. Al-Wahaibi, and A.H. Muggeridge. 2008. Physical porperties (densisy, excess

molar volume, viscosity, surface tension and refractive index) of ethanol + glycerol. J. Chem. Eng. Data. 53: 2793–2796.

2. Williams, A.T.R., S.A. Winfield, and J.N. Miller. 1983. Relative fluorescence quantum yields using a computer-controlled luminescence spectrometer. Analyst. 108: 1067.

3. Gullapalli, R.R., T. Tabouillot, R. Mathura, J.H. Dangaria, and P.J. Butler. 2015. Integrated multimodal microscopy, time-resolved fluorescence, and optical-trap rheometry: toward single molecule mechanobiology. J. Biomed. Opt. 12: 014012.

4. Penzkofer, a, and Y. Lu. 1986. Fluorescence Quenching of Rhodamine 6G in Methanol At High Concentration. Chem. Phys. 103: 399–405.

5. Olson, F., C.A. Hunt, F.C. Szoka, W.J. Vail, and D. Papahadjopoulos. 1979. Preparation of liposomes of defined size distribution by extrusion through polycarbonate membranes. Biochim. Biophys. Acta - Biomembr. 557: 9–23.

6. Galbraith, C.G., K.M. Yamada, and M.P. Sheetz. 2002. The relationship between force and focal complex development. J. Cell Biol. 159: 695–705.

7. Levitus, M., and S. Ranjit. 2011. Cyanine dyes in biophysical research: the photophysics of polymethine fluorescent dyes in biomolecular environments. Q. Rev. Biophys. 44: 123–151.

8. Muddana, H.S., S. Sengupta, A. Sen, and P.J. Butler. 2014. Enhanced brightness and photostability of cyanine dyes by supramolecular containment. arXiv:1410.0844. : 1–20.

9. Widengren, J., and P. Schwille. 2000. Characterization of Photoinduced Isomerization and Back-Isomerization of the Cyanine Dye Cy5 by Fluorescence Correlation Spectroscopy. : 6416–6428.

10. Aramendia, P.F., R.M. Negri, and E.S. Roman. 1994. Temperature Dependence of Fluorescence and Photoisomerization in Symmetric Carbocyanines.Influence of Medium Viscosity and Molecular Structure. J. Phys. Chem. 98: 3165–3173.

11. Loutfy, R.O., and B.A. Arnold. 1982. Effect of viscosity and temperature on torsional relaxation of molecular rotors. J. Phys. Chem. 86: 4205–4211.

12. Doolittle, A.K. 1952. Studies in newtonian flow. III. the dependence of the viscosity of liquids on

molecular weight and free space (in homologous series). J. Appl. Phys. 23: 236–239.

13. Li, X.M., J.M. Smaby, M.M. Momsen, H.L. Brockman, and R.E. Brown. 2000. Sphingomyelin interfacial behavior: the impact of changing acyl chain composition. Biophys. J. 78: 1921–31.

14. Petrache, H.I., S.W. Dodd, and M.F. Brown. 2000. Area per Lipid and Acyl Length Distributions in Fluid Phosphatidylcholines Determined by 2H NMR Spectroscopy. Biophys. J. 79: 3172–3192.

15. Kucerka, N., Y. Liu, N. Chu, H.I. Petrache, S. Tristram-Nagle, and J.F. Nagle. 2005. Structure of Fully Hydrated Fluid Phase DMPC and DLPC Lipid Bilayers Using X-Ray Scattering from Oriented Multilamellar Arrays and from Unilamellar Vesicles. Biophys. J. 88: 2626–37.

16. Kucerka, N., M. a Kiselev, and P. Balgavý. 2004. Determination of bilayer thickness and lipid surface area in unilamellar dimyristoylphosphatidylcholine vesicles from small-angle neutron scattering curves: a comparison of evaluation methods. Eur. Biophys. J. 33: 328–334.

17. Nagle, J.F., and S. Tristram-Nagle. 2000. Structure of lipid bilayers. Biochim Biophys Acta. 1469: 159–195.

18. W.-J. Sun, R. M. Suter, M. A. Knewtson, C. R. Worthington, S. Tristram-Nagle, R. Zhang, and J.F.N. 1994. Order and disorder in fully hydrated unoriented bilayers of gel-phase dipalmitoylphosphatidylcholine. Phys. Rev. E. 49: 4665–4676.

19. Sun, W.J., S. Tristram-Nagle, R.M. Suter, and J.F. Nagle. 1996. Structure of gel phase saturated lecithin bilayers: temperature and chain length dependence. Biophys. J. 71: 885–891.

20. Wiener, M.C., and S.H. White. 1992. Structure of a fluid dioleoylphosphatidylcholine bilayer determined by joint refinement of x-ray and neutron diffraction data. II. Distribution and packing of terminal methyl group. Biophys. J. 61: 428–433.

21. Kucerka, N., J.J.F. Nagle, J.N.J. Sachs, S.E.S. Feller, J. Pencer, A. Jackson, J. Katsaras, and N. Kučerka. 2008. Supplement - Lipid bilayer structure determined by the simultaneous analysis of neutron and X-ray scattering data. Biophys. J. 95: 2007.

22. Smaby, J.M., M.M. Momsen, H.L. Brockman, and R.E. Brown. 1997. Phosphatidylcholine acyl unsaturation modulates the decrease in interfacial elasticity induced by cholesterol. Biophys. J. 73:

1492–505.

23. Nagle, J.F. 2013. Introductory Lecture: Basic quantities in model biomembranes. Faraday Discuss. 161: 11–150.

24. Himbert, S., R.J. Alsop, M. Rose, L. Hertz, A. Dhaliwal, J.M. Moran-Mirabal, C.P. Verschoor, D.M.E. Bowdish, L. Kaestner, C. Wagner, and M.C. Rheinstädter. 2017. The Molecular Structure of Human Red Blood Cell Membranes from Highly Oriented, Solid Supported Multi-Lamellar Membranes. Sci. Rep. 7: 39661.

25. Pan, J., S. Tristram-Nagle, N. Kučerka, and J.F. Nagle. 2008. Temperature Dependence of Structure, Bending Rigidity, and Bilayer Interactions of Dioleoylphosphatidylcholine Bilayers. Biophys. J. 94: 117–124.
